# Initial and corrective submovement encoding differences within primary motor cortex during precision reaching

**DOI:** 10.1101/2023.07.01.547340

**Authors:** Kevin C Schwartze, Wei-Hsien Lee, Adam G Rouse

## Abstract

Precision reaching tasks often require corrective submovements for successful completion. Most studies of reaching have focused on single initial movements, and the cortical encoding model was implied to be the same for all submovements. However, corrective submovements may show different encoding patterns from the initial submovement with distinct patterns of activation across the population. Two rhesus macaques performed a precision center-out-task with small targets. Neural activity from single units in primary motor cortex and associated behavioral data were recorded to evaluate movement characteristics. Neural population data and individual neuronal firing rates identified with a peak finding algorithm to identify peaks in hand speed were examined for encoding differences between initial and corrective submovements. Individual neurons were fitted with a regression model that included the reach vector, position, and speed to predict firing rate. For both initial and corrective submovements, the largest effect remained movement direction. We observed a large subset changed their preferred direction greater than 45° between initial and corrective submovements. Neuronal depth of modulation also showed considerable variation when adjusted for movement speed. By utilizing principal component analysis, neural trajectories of initial and corrective submovements progressed through different neural subspaces. These findings all suggest that different neural encoding patterns exist for initial and corrective submovements within the cortex. We hypothesize that this variation in how neurons change to encode small, corrective submovements might allow for a larger portion of the neural space being used to encode a greater range of movements with varying amplitudes and levels of precision.

**New and Noteworthy:** Neuronal recordings matched with kinematic behavior were collected in a precision center-out task that often required corrective movements. We reveal large differences in preferred direction and depth of modulation between initial and corrective submovements across the neural population. We then present a model of the neural population describing how these shifts in tuning create different subspaces for signaling initial and corrective movements likely to improve motor precision.

## Introduction

Our brain allows us to execute movement over a large dynamic range. While large, ballistic movements typically need less fine control, precision movements require it (Abrams et al. 1990; Craik 1948; Elliott et al. 2010; Sainburg et al. 1999; Woodworth 1899). One reaching movement can be broken down into 2 types of submovements: an initial submovement or any following corrective submovements (Milner and Ijaz 1990; Novak et al. 2002; Port et al. 1997; Pratt et al. 1994). Reaching movements have bell-shaped velocity profiles (Flash and Hogan 1985; Georgopoulos et al. 1981; Morasso 1981; Soechting 1984), a feature seen in both the initial and corrective submovement components (Hatsopoulos et al. 2007). However, by their nature, corrective submovements have a significantly smaller magnitude than the initial submovement.

The firing rates of individual neurons in the motor cortex signal information about upcoming movements. Populations of individual neurons within the motor cortex represent several reaching characteristics of action selection and execution (Kalaska et al. 1997), including force generation (Evarts 1968) and preferred direction (Fu et al. 1993; Georgopoulos et al. 1982, 1986). Technical advances have allowed for recording from more neurons to sample and build accurate predictions of upcoming movement features from specific neuronal firing within the cortex, (Kalaska and Crammond 1992; Moran and Schwartz 1999; Nicolelis et al. 1997). Models can describe how neural activity depends on various movement conditions (Moran and Schwartz 1999; Wang et al. 2007). Condition dependent activity is the modulation of neural firing represented by movement specific parameters such as direction, speed, and spatial location (Georgopoulos et al. 1986; Moran and Schwartz 1999; Rouse, Adam G., Schieber 2017; Rouse and Schieber 2016; Wang et al. 2007).

Most experiments of neural encoding for reaching have only examined instructed movements that were generally large, overlearned, and stereotyped to allow for trial averaging. For some analyses and applications such as brain-computer interfaces (BCIs), identical models of encoding have been assumed to fit both initial and corrective submovements. However, we hypothesize that initial and corrective submovements are encoded differently (Evarts and Fromm 1977; Fu et al. 1995; Milner 1992; Scott 2004), allowing for a greater diversity of successive submovements. Understanding the fundamental differences between submovement planning, execution, and implementation is vitally important for real-world applications such as brain-computer interface algorithms (Umair et al. 2017). Current BCI prostheses are capable of performing basic functions; however, users struggle to complete precision tasks with the devices (Mridha et al. 2021). Recent BCI algorithms have started incorporating increasingly complex decoding systems (Inoue et al. 2018; Karpowicz et al. 2022; Ma et al. 2022), but more information about the underlying organization in the brain of corrective movements in healthy reaching will help drive further development.

As researchers record larger populations of motor cortex neurons during reaching movements, the results show previously indiscernible elegant patterns of neural activity (Churchland et al. 2006; Churchland and Shenoy 2007; Maynard et al. 1999). These neural firing patterns can be temporally mapped in multidimensional neural subspaces as (Churchland et al. 2012; Kaufman and Churchland 2016; Rouse and Schieber 2018) using dimensionality-reduction tools, such as principal components analysis (PCA). Comparison of neural trajectories between initial and corrective submovements is thus far unquantified and would provide important information about cortical encoding patterns during precision reaching.

In the current study we utilized a custom precision center-out task to encourage multi-phasic submovement corrections. We previously showed that condition-invariant neural activity reliably defines the timing of both the initial and corrective submovements (Rouse et al. 2022). We now ask how neural firing rates change with the specific reach conditions of velocity, position, and speed during initial and corrective submovements. Kinematic behaviors were aligned with primary motor cortex neural recordings to evaluate encoding pattern differences between initial and corrective submovements. We utilized a well-established cortical tuning model (Wang et al. 2007) to evaluate encoding differences between submovement types as well as modeled the observed trajectories of the observed neural tuning to measure shared encoding subspaces.

## Methods

### Macaque Model

Two adult, male rhesus macaques (Macaca mulatta), P and Q, were used in the research project. All standard procedures for the care and use of these nonhuman primates followed the Guide for the Care and Use of Laboratory Animals and received approval from the University Committee on Animal Resources at the University of Rochester Medical Center, Rochester, NY.

### Experimental Design

Subjects P and Q were trained to perform a precision center-out-task using a custom interface equipped with a joystick and monitor (Figure 1A). The joystick was 18 cm and had the centering spring removed, allowing it to move freely with little resistance. The precision task required the subject to single handedly manipulate the joystick moving the displayed cursor from the center into an indicated goal area. The goal areas were spaced equidistant from each other in 45° increments, and designed to be narrow and deep, wide and shallow, or wide and deep (Figure 1B).

**Figure 1.**
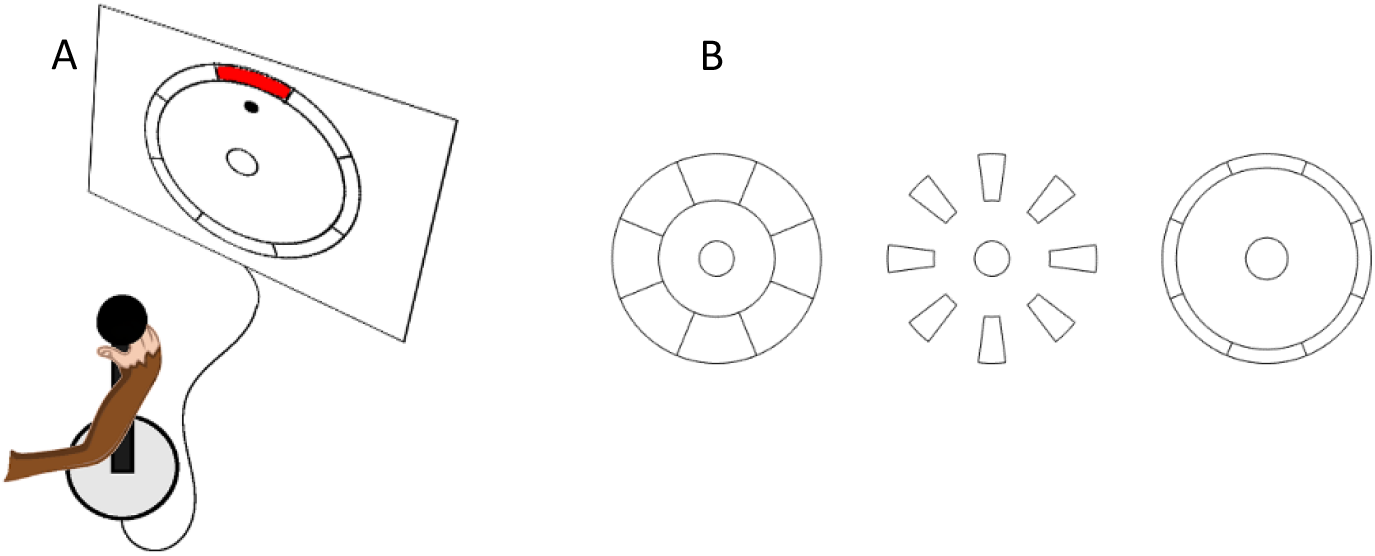
Precision center-out-task design. A) Rhesus macaques manipulated a joystick controlling the cursor on a digital screen. The cursor was required to move from the center starting area to the indicated peripheral target zone. B) Peripheral target zone layouts designed to encourage corrective submovements during precision reaching. Target zones are wide and deep (left), narrow and deep (middle), or wide and shallow (right).

Our precision center-out-task utilized small targets and a required hold time to elicit not only initial center-out reaches but also corrective movements. After reaching the goal area, the animals were required to hold for 500-600 ms. If the cursor left the goal area within the hold interval, we allowed them to move and re-enter the goal as many times as required to complete a successful hold. Animals were provided a liquid reward upon successful trial completion. Additional animal and task details are described in Rouse et al. 2022.

### Neural Recordings

Both subjects P and Q were surgically implanted with chronic multielectrode arrays to detect individual neuronal activity. Floating microelectrode arrays (MicroProbes for Life Science) were implanted in primary motor cortex (M1) in each monkey, using methods described in detail previously (Mollazadeh et al. 2011; Rouse et al. 2022; Rouse and Schieber 2016). For monkey P, recordings were collected from six 16-channel arrays implanted in M1. For monkey Q, two 32-channel arrays and one 16-channel array in M1 were used. Channels with spiking activity were collected with Plexon MAP (Plexon) and Cerebus (Blackrock Microsystems, LLC.), thresholded manually online, and sorted off-line with a custom, semi-automated algorithm.

### Behavior Analysis

Our previous work found that most movements in our precision center-out-task are well defined as distinct submovements with bell-shaped speed profiles separated by low-speed intervals. Here, we again identified initial and corrective submovements by identifying cursor speed peaks (Figure 2) using a peak finding algorithm as previously described in Rouse et al (2022). Briefly, speed peaks were defined as having a i) speed >250 pixels/s, ii) no larger peak within ±200-ms, and iii) having a peak prominence of at least 50% of its two adjacent speed troughs. All speed-peak defined submovements from leaving the center until completion of a successful hold were identified. The initial submovement was defined as the large initial reach that moved at least halfway to the goal area and all subsequent submovements were defined as corrective. Rare small amplitude submovements before the initial submovements as well as submovements made entirely within the goal area were discarded from analysis.

**Figure 2.**
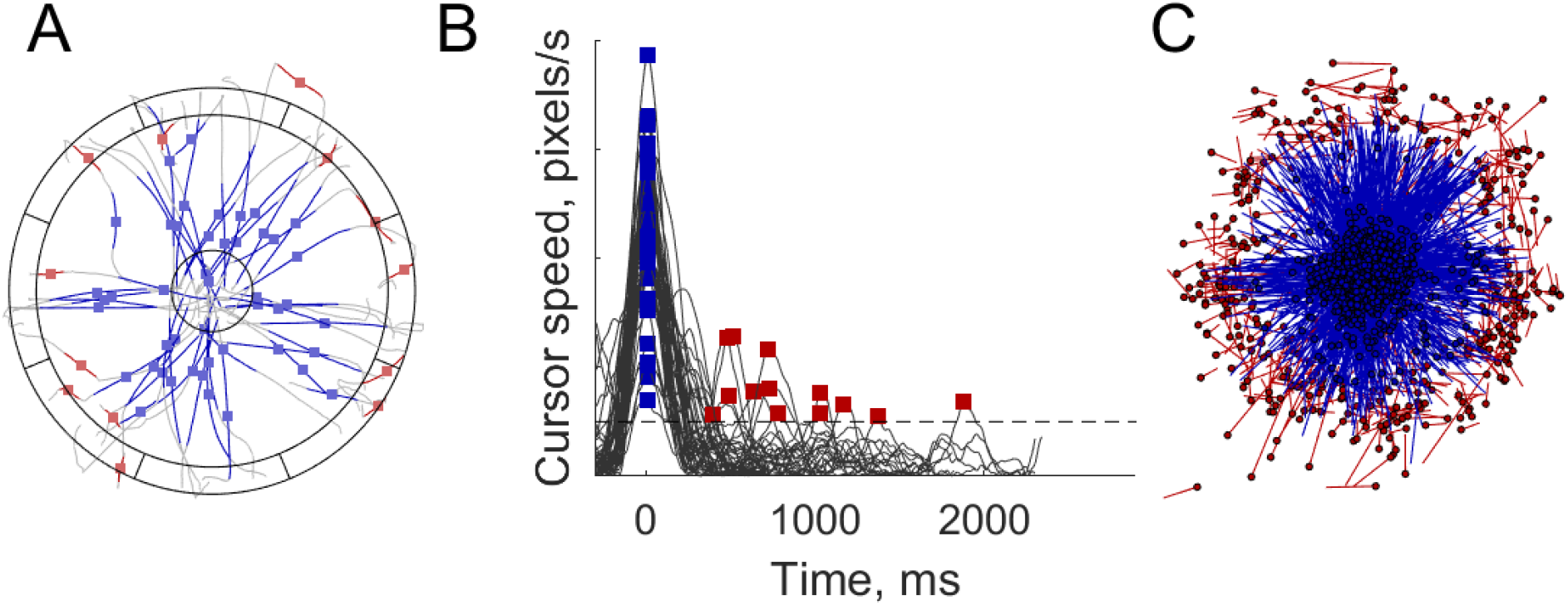
Submovement identification and vector approximations. A) Movements performed on the wide and shallow peripheral target. Initial submovements are traced in blue and corrective submovements are in red with speed peak occurrence noted with a circle. B) Speed profiles for the submovements in (2A) with initial speed peaks denoted with a blue square and corrective speed peaks noted with a red square. The dotted line shows the 250pixel/s threshold for submovement designation. C) Initial (blue) and corrective (red) submovements approximated as single line vectors. Submovement starting location indicated with a circle.

### Movement Vectors

All submovements, both initial and corrective, were approximated as straight-line vectors using the beginning and end cursor locations (Figure 2C) to simplify the analysis of the relationship between neural activity and submovement kinematics. Each vector is comprised of a starting position, the movement direction, and distance travelled within the 200 ms time window from 100 ms before peak speed until 100 ms after. All initial submovements had a similar starting point in the center region, as expected during a center-out task. Most corrective submovements started in the periphery of the workspace near the goal areas.

### Multivariate Regression

Multivariate linear regression (Figure 3) was performed on initial and corrective submovements to analyze correlations of the kinematics to neural firing rates. We used the established cortical activity model published by Wang et al. (2007). This model included 3 features: position, velocity, and speed defined by the straight-line vector. Position for each submovement was defined as the starting movement position 100 ms before peak speed. Velocity was the submovement vector from movement onset (100 ms before) to offset (100 ms after). Speed was the magnitude of the submovement vector.

**Figure 3.**
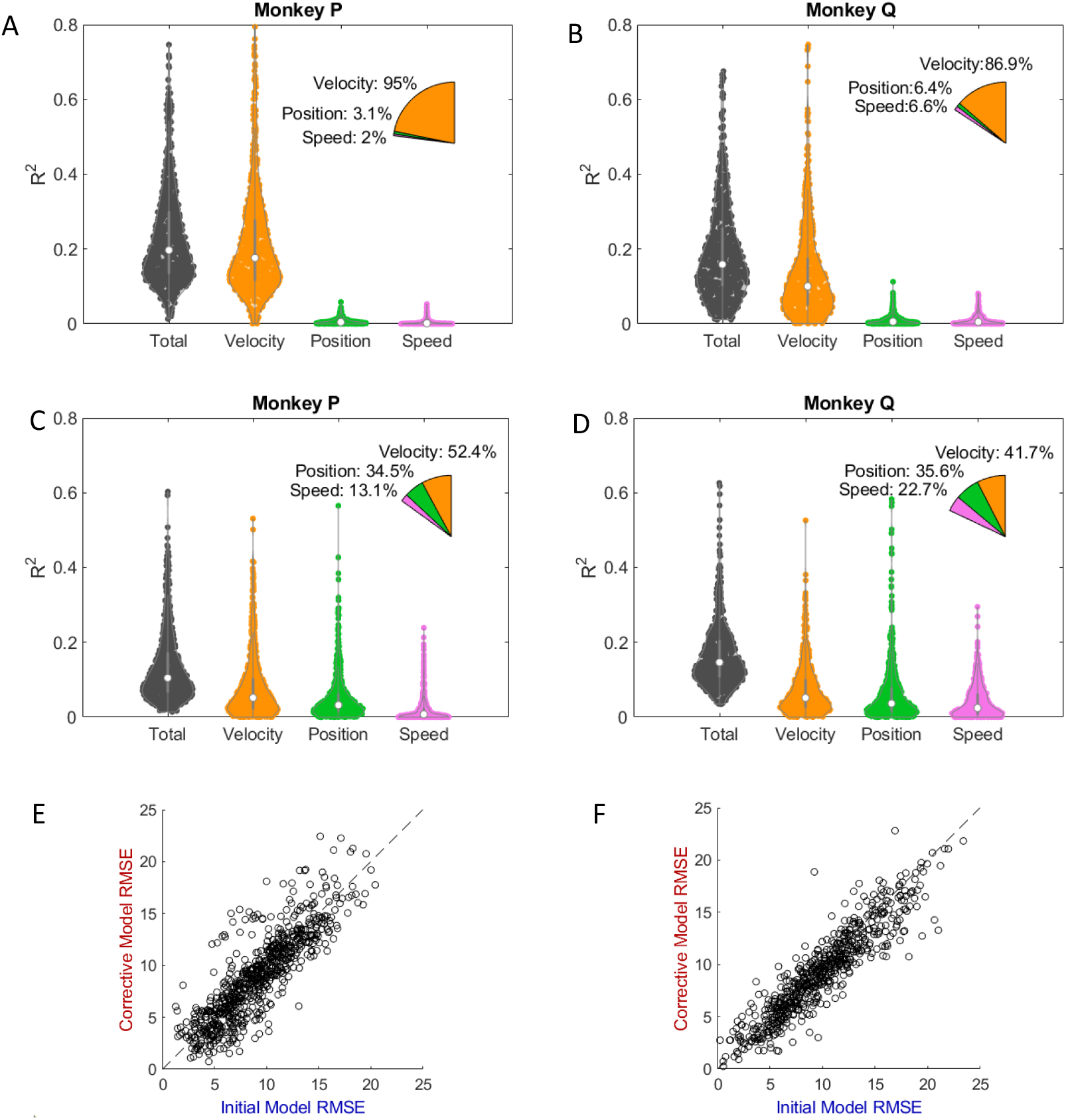
Multivariate regression analysis. A, B) Partial R^2^ for firing rate explained for the velocity vector, starting hand position, and movement speed of all tuned neurons using multivariate regression for initial and C, D) corrective submovements for monkeys P and Q. Violin plots courtesy of Bechtold (Bechtold 2016). Partial pie charts represent the overall the regression elements in proportion to each other and the percent contribution to the total. E, F) Root-mean-square-error (RMSE) analysis of initial and corrective submovements for monkeys P and Q. The dotted line shows equivalence for residual noise between initial and corrective. Correlation values for E) monkey P and F) monkey Q are 0.85 (0.83-0.87, 95% confidence interval) and 0.92 (0.91-0.93, 95% confidence interval), respectively.

The regression model uses the following equation:

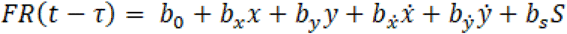

Here *x* and *y* are the starting position such that *x*=0 and *y*=0 defines the center. The velocity vectors are defined by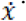 and 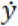. The speed, *S*, was the magnitude of the velocity vector.

Firing rates were calculated with a Gaussian smoothing window (σ=30 ms) and sampled in 10ms bins. A sliding time lag (τ) from 300 ms before until 100 ms after peak movement speed was used to identify and select the highest total R^2^ for each spiking unit for initial and corrective submovements separately. Once this single time lag was identified, the spiking units’ relation to kinematic features was examined. Since each session had 600-1100 trials, the p-values for the multivariate regression were likely to be statistically significant with the large number of datapoints even if the effect was small. We instead chose to include spiking units based on effect size using the fraction total variance explained as measured with R^2^. We used an R^2^ greater than 0.1 for defining the spiking units that meaningfully modulated with the task and included units that had an R^2^ greater than 0.1 for either initial or corrective submovements.

The partial R^2^ describes how much a spiking unit’s variance in firing rate is attributable to each predictor. The partial R^2^ is calculated by the increase in variance explained by the full compared to a model without the given predictor (Pardoe 2020). We calculated the partial R^2^ for starting position (*x, y*), velocity 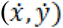, *and speed (S)*. (For the velocity partial R^2^ we used the following equation:

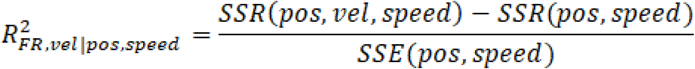

The partial R^2^ for velocity is the regression sum of squares (SSR) of the full regression including velocity along with position and speed minus the reduced model regression sum of squares of only position and speed divided by the sum of square errors of the reduced model. The position and speed partial R^2^ values were calculated analogously comparing the full to reduced model.

We were primarily interested in how the velocity tuning changed between initial and corrective submovements after correcting for differences in starting position and speed. A neuron’s preferred reach direction (θ_Pref_) is the direction with maximal firing rate and can be defined as:

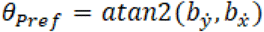

Using the 2-argument arctangent of the x and y velocity coefficients, 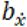 and 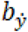.

A neuron’s depth of modulation (DOM) measures how much the firing rate changes between reaches in the preferred and anti-preferred direction.

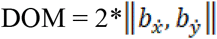

It is the combined magnitude of the x and y velocity coefficients, 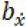 and 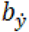 and doubled to account for the change above and below baseline for the preferred and anti-preferred direction. We were interested in both the actual change in firing rates that was observed for initial and corrective submovements during the experimental task as well as the depth of modulation as a function of velocity. We refer to the change in firing rate between the preferred and anti-preferred direction observed for an average magnitude submovement (either initial or corrective) during the experiment as the task depth of modulation. We then also calculated the depth of modulation scaled for the velocity of initial and corrective submovements. We refer to this as the velocity depth of modulation which provides a common scale to compare the depth of modulation relative to velocity across both initial and corrective submovements.

For examining the relationship between firing rate and reach direction, we display the marginal firing rates corrected for speed and starting position when firing rates as a function of reach direction are shown.

To estimate the confidence interval (C.I.) for the preferred direction estimate, bootstrapping by drawing with replacement over 10,000 iterations from the spiking unit population deemed to be significantly tuned. The 95 percent C.I. was then taken to be the range between the 2.5 and 92.5^th^ percentile of the bootstrapped samples.

### Neural Population Model

A neural population model was built using the cortical tuning parameters identified with our multivariate regression to further explore the differences in neural tuning between initial and corrective submovements. Since the corrective submovements were idiosyncratic due to motor error and not repeated to allow for trial averaging, we instead simulated the neural activity for initial and corrective submovements to each of the 8 center-out directions. These simulated movements were assumed to be all the same magnitude as the average speed of either initial or corrective submovements. The linear tuning parameters for each spiking unit as a function of time were then used to generate simulated neural trajectories to the 8 different targets.

Principal component analysis was used to identify the 2-dimensional neural subspace with the most variance at the timepoint 150 ms before peak speed for both initial and corrective submovements. These subspaces were calculated for both initial and corrective neural activity and the opposite neural activity was also projected into the separately identified subspaces. The percentage of total variance accounted for simulated activity by these subspaces for all activity from 300 ms before until 100 ms after peak speed was calculated. Additionally, the total overlap of simulated neural activity (Elsayed et al. 2016; Rouse and Schieber 2018) was calculated using the formula:

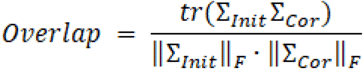

The 2 covariance matrices for initial and corrective neural activity (*Σ*_*Init*_, *Σ*_*Cor*_) are compared by using the trace of the multiplied covariance matrices to calculate the overlap. This result is normalized by the Frobenius norm (*‖ … ‖*_*F*_) of each covariance matrix to obtain a metric between 0 (completely orthogonal neural activity) and 1 (complete alignment).

## Results

### Multivariate Regression

Subjects P and Q successfully completed 10964 and 8126 trials, respectively. With these trials, 6007 required at least one identified corrective submovement with a total of 10391 corrective submovements across all trials. Individual neurons from each session were fit with a multivariate linear regression model separately for all initial and corrective submovements. It included the 2-dimensional reach direction vector, hand position, and hand speed to predict firing rate. Neurons were considered tuned with an R^2^ value of greater than 0.1 for either initial or corrective submovements. A total of 1293 tuned neurons were identified for subject P and 1185 for Subject Q. For initial submovements, the largest effect for almost all units was velocity (Figure 3A,B). This was expected as there was little variability of speed or position for the initial submovements as the starting hand position and distance to target remained approximately the same for all trials. For corrective submovements analysis, the influence of the velocity vector remained the largest factor; however, it decreased overall when compared to initial submovements (Figure 3C,D). Significantly larger impacts by hand position and speed were both observed, with hand position having a higher R^2^ in both subjects. We examined the residual noise with the root-mean square error (RMSE), for each unit after the fitting the three factors of velocity, position, and speed. A similar amount of deviation for individual reaches was observed. Suggesting a relatively similar ability of our multivariate regression to predict neural activity in both initial and corrective submovements. Albeit with a lower R^2^ due to the generally decreased modulation of firing rate for corrective movements.

### Preferred Direction

We next used the linear cortical tuning model for each unit to generate its cosine tuning curve as a function of reach direction. The direction with the largest increase of firing rate is defined as the preferred direction with the direction 180° opposite being the anti-preferred direction. We analyzed the preferred direction of each spiking unit for both initial and corrective submovements separately. If both submovements are encoded identically, the modeled preferred direction would be the same. For both subjects, the majority of neurons maintained approximately the same directional tuning, with less than a 45° change in preferred direction between submovement types (Figure 4B,C); however, a significant number of neurons demonstrated a change in preferred direction to an entirely different goal target location. Subject P had 42.5% (39.0-46.0%, 95% C.I.) of tuned neurons change their preferred direction at least 45° between submovement types (Figure 4B). Subject Q had a similar representation of 40.0% (36.1-43.7%, 95% C.I.) of neurons changing preferred direction (Figure 4C).

**Figure 4.**
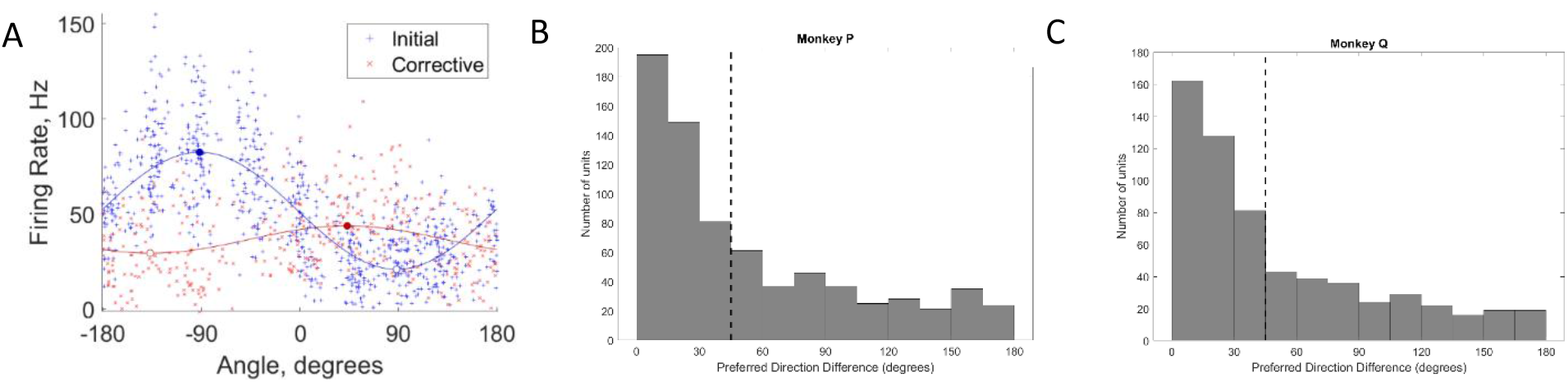
Preferred direction analysis for all tuned (R^2^>0.1) M1 neurons. A) The cosine tuning curve for an example neuron with both initial (blue) and corrective (red) submovements. The preferred (max firing rate per direction) directions were −92° for initial and 43° for corrective submovements (marked with filled circles). Note, the firing rate may be slightly negative for some submovements due to the correction made for position and speed to show the relationship to reach direction. B) The number of neurons with a ≥45° change of preferred direction encoding between initial and corrective submovements for Subject P and C) Subject Q.

### Depth of Modulation

The task depth of modulation is the firing rate change between the preferred and anti-preferred directions (Figure 5A). As expected for most spiking units, a larger depth of modulation was observed during initial submovements than during corrective submovements during the task (5B,C). For a single linear encoding model, proportional increases in modulation would be expected to accompany increases in movement speed. However, when the depth of modulation is scaled linearly for velocity, the changes in firing rate showed considerable variation (Figure 5D,E). While some neurons exhibited the changes along the 45° diagonal consistent with a 1x fixed linear relationship with speed, some neurons were primarily active only during initial submovements (<1x corrective to initial DOM ratio) and other neurons exhibited significantly increased gain in firing rate for the small corrective submovements (>1x), and. No clear relationships were observed suggesting different patterns of neural activity at the individual neuron level for the two submovement types.

**Figure 5.**
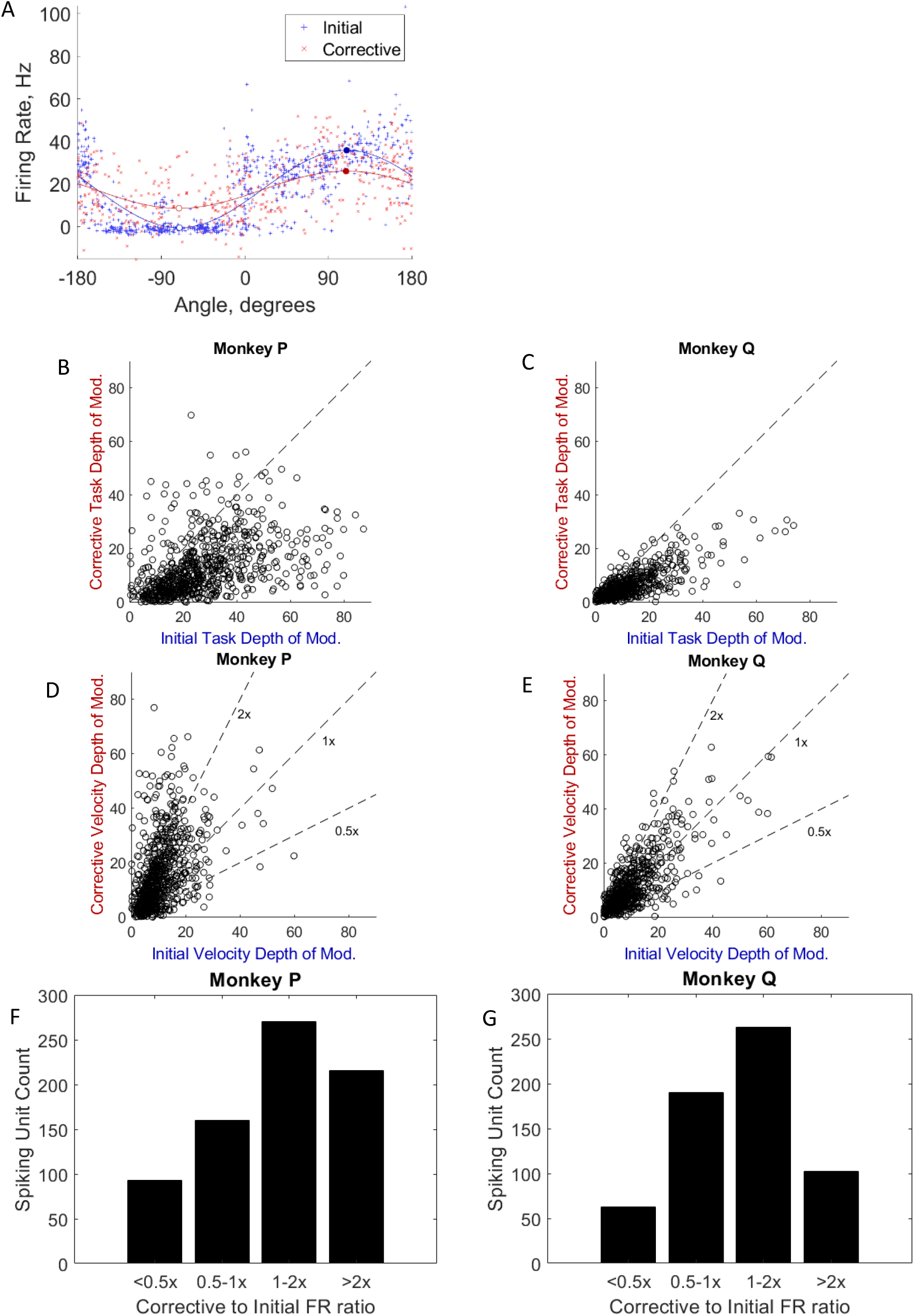
Depth of modulation analysis for tuned M1 neurons. A) The cosine tuning curves of an example neuron for initial (blue) and corrective (red) submovements. The task depth of modulation is the difference in firing rate between the preferred and anti-preferred directions for each submovement type (bold blue and red vertical lines). The depth of modulation was 36.3 Hz for initial and 17.4 Hz for corrective for this example neuron. Note, the firing rate may be slightly negative for some submovements due to the correction made for position and speed to show the relationship to reach direction. B, C) Direct comparison of all individual neurons’ task depth of modulation analysis for all R^2^>0.1 units for initial and corrective submovements for monkey P and monkey Q. The dotted line shows equivalence task DOM for both initial and corrective. D, E) Velocity depth of modulation comparison for all R^2^>0.1 units for monkey P and monkey Q. The dotted lines show 2x, 1x, and 0.5x corrective to initial velocity DOM ratios. F) Velocity DOM for spiking units separated by activity dedicated to primarily initial or corrective submovements for monkey P and G) monkey Q. Correlation and confidence intervals, respectively, for F) monkey P of 0.47 and 0.41-0.53 and for G) monkey Q of 0.75 and 0.72-0.79.

### Neural population model

To further explore the tuning observed across the neural population, we built a simulated neural population with the tuning parameters observed. Using each spiking unit’s tuning coefficients for initial and corrective submovements, we generated simulated neural activity for an entire population as if the subject performed the same amplitude center-out reaches in 8 directions. One set of firing rates simulated the tuning observed for initial and one set for corrective submovements. We next used principal component analysis (PCA) to separately identify and display the neural subspaces with the most initial and corrective variance. We then plotted the neural trajectories of the modeled firing rates for both initial and corrective submovements in these separate subspaces. In the initial subspace, larger magnitude neural activity occurred for the initial neural data than the corrective, as expected (top row of Fig. 6). In the separate corrective subspace, however, larger firing rate changes occurred in these dimensions for the corrective neural data (bottom row of Fig. 6). Suggesting better representation of these corrective, smaller amplitude movements in different neural dimensions. The shifts in tuning indicate neural dimensions with increased neural activity depending on execution of an initial or corrective submovement. We hypothesize that different neural dimensions—as created by shifts in preferred direction and depths of modulation for a subset of neurons—allow for more accurate encoding of small precise movements than if the same, global neural subspace signaled both initial and corrective movements.

**Figure 6.**
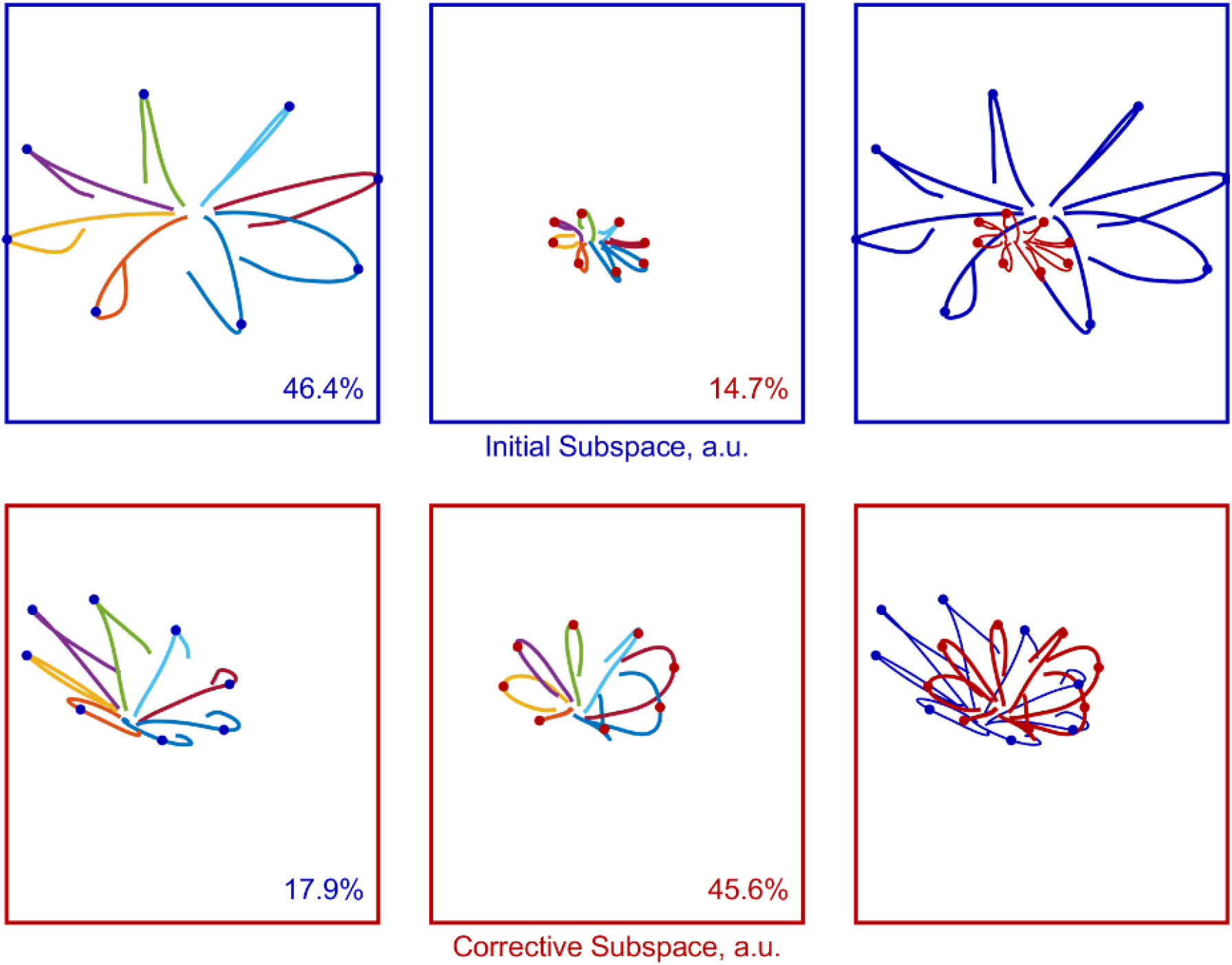
Modeled firing rates using tuning parameters observed during initial and corrective submovements. Two different neural subspaces maximized variance for initial (top) and corrective (bottom) submovements. Left) The neural activity for simulated reaches with tuning observed for initial submovements. Middle) The neural activity for simulated reaches with tuning observed for corrective submovements. Right) Initial (blue) and corrective (red) neural tuning shown in the initial and corrective subspaces. The colors (left and middle) represent reaches to the 8 center-out targets. The dots indicate neural activity 150 ms before peak speed. The shifts in tuning indicate neural dimensions with increased neural activity depending on execution of an initial or corrective submovement. Percentages indicate how much of the total neural activity is accounted for in the given two-dimensional subspace. For monkey Q (neural trajectory plot not shown), the two-dimensional subspace maximized for initial and corrective neural activity accounted for 52.8% and 47.1% of total neural activity, respectively. The initial subspace accounted for 33.1% of corrective and the corrective subspace accounted for 35.7% of initial. The total overlap in simulated neural activity between initial and corrective submovements was 0.31for monkey P and 0.45 for monkey Q.

## Discussion

Utilizing a custom precision center-out task, we quantified differences of several neural tuning parameters between initial and corrective submovements within the primary motor cortex. While differences are not unexpected, quantifying these differences is an important step in describing neural tuning to not only instructed movements but also subsequent corrective movements and could be a key foundation for BCI encoders to build upon. Our analysis primarily focused on individual neuron’s preferred direction and depth of modulation.

### Shifts in Neural Tuning

Directional tuning of neurons in the primary motor cortex is well established (Fu et al. 1993; Georgopoulos et al. 1982, 1986). We have demonstrated that a significant portion of individual neurons in primary motor cortex change their preferred direction when encoding for corrective versus initial submovements. We observed approximately 40% of spiking units were classified to have a >45° shift in preferred direction, an entire target area shift in the task protocol.

It has been well documented that larger and faster movements are typically elicited by increased neural descending signals (Desmedt and Godaux 1977; Farina et al. 2002; Freund 1983; Milner-Brown et al. 1973). As expected, our results showed an overall higher firing rate towards initial submovements, as would be expected due to their innately larger magnitude; however, when we adjusted for movement velocity, a range of relationships between speed and individual cells became apparent. Some neurons fired almost exclusively during initial submovements, and others exhibited a much larger response for corrective submovements than expected for a linear speed relationship. Rather than uniform increases in firing rate, our results align with sequential recruitment of some new cortical neurons with increasing speeds similar to the Henneman’s size principal for motoneurons (1965). Additionally, a small fraction of units are even more active for smaller movements than larger (Evarts et al. 1983), highlighting that neural tuning to speed is far from consistent across the neural population.

### Different Neural Subspaces

The observed shifts and modulation in neural tuning lead us to propose that different neural subspaces are utilized within the motor cortex to encode the initial and corrective submovements. These different neural subspaces can be thought of as different combinations or patterns of neurons selectively activated for movements in the same direction but with different amplitudes or task demands. The many neurons with shared preferred directions between initial and corrective submovements likely are due to the same agonist and antagonist muscles being responsible for reaching in each direction. However, for different magnitude movements, the amount of co-contraction and balance between agonist/antagonist muscle may shift along with changes in fixator muscle demands (Griffin et al. 2015).

Beyond the muscle activation differences, different subspaces potentially provide an elegant means to adjust the signal gain between cortical firing rates and muscle activation levels (Elsayed et al. 2016). The larger modulation in firing rate within a separate neural subspace mapped to small amplitude movements could provide more accurate encoding of small, precise movements compared to a system that uses only a single subspace with smaller changes in firing rate. If we assume cortical neurons and corresponding subspaces may be constrained by similar levels of noise, the larger signal mapping to smaller motor output would have a better signal-to-noise ratio. A subspace best suited for precision movement execution while other subspaces may be better for faster movements or other functions. Thus, expanding the motor capabilities without increasing the number of neurons required. Further studies are needed to better define and verify these diverse and dynamic encoding patterns undefinable by a single cortical tuning model.

The relationship between movement amplitude and target size in Fitt’s Law is well established (Fitts 1964; MacKenzie and Buxton 1992; Monk and Jagacinski 1985; Mottet et al. 1994; Murata 1999; Paul M. Fitts 1954; Takeda et al. 2019); however, the neural mechanisms that underlie Fitt’s Law remain elusive. A single, linear cortical model is unable to reproduce the motor control necessary for increased precision with smaller movements. Varying neural dynamics allowed by multiple neural subspaces might help optimize movement parameters (direction, speed, etc.) to accomplish a given task. Additionally, the increased sensory integration with movement execution for a precision task are likely to alter the underlying neural mechanisms encoding movement. Better defining these neural states may lead to better descriptions of the dynamic response required for precision movements that lead to Fitt’s Law.

### Limitations

By relying on user-generated errors, we were not able to systematically vary which targets required corrective submovements. Additionally, with the center out task, there was greater variety of starting position for corrective submovements than for initial. This is represented in the multivariate regression analysis (Figure 3) with the larger position effect on firing rates for corrective submovements. Despite this added influence, the velocity vector remained the primary factor for neural firing rates for both types of submovements lending credence to velocity and not position being the dominantly encoded motor cortex feature.

Noise in the model fits could explain part of the observed shift in neural tuning. In a direct comparison of root-mean-square-error (RMSE) of neural firing rates between initial and corrective submovements, we saw similar noise variance for predictions of both types of submovements (Figure 3E, F). Thus, it appears the model error due to noise was similar for both the initial and corrective data and strengthens our conclusions that true changes in preferred direction and depth of modulation are occurring for initial and corrective submovements (Stein et al. 2005).

Further experiments with all combinations of large and small movements that are both instructed and corrective are needed to more completely quantify these shifts in tuning. In future studies tasks that produce small initial submovements as well as small natural corrective submovements should be explored. Instructed corrective submovements induced by a change of target location or external force affecting arm trajectory should also be explored.

## Conclusion

In summary, we have quantified the shifts in neuronal tuning within the primary motor cortex for initial versus corrective submovements using a precision reaching center-out-task. These shifts resulted in poor description of both submovement types by a single fixed linear model. We have highlighted how these shifts when viewed from a neural space perspective give evidence of different neural subspaces being utilized for generation of initial and corrective submovements.

## Acknowledgements

We would like to thank Dr. Marc Schieber and his laboratory at the University of Rochester Medical Center for assistance with data collection.

## Data Availability

Complete dataset is available at https://doi.org/10.6084/m9.figshare.23631951

Analysis results are available at https://doi.org/10.6084/m9.figshare.23620236

Analysis code available at https://zenodo.org/badge/latestdoi/660266065

## References

Abrams RA, Meyer DE, Kornblum S. Eye-hand coordination: oculomotor control in rapid aimed limb movements. J Exp Psychol Hum Percept Perform 16: 248–267, 1990.

Bechtold B. Violin plots for matlab [Online]. Github Proj. 2016.https://github.com/bastibe/Violinplot-Matlab, DOI: 10.5281/zenodo.4559847.

Churchland MM, Cunningham JP, Kaufman MT, Foster JD, Nuyujukian P, Ryu S, Shenoy K V. Neural population dynamics during reaching. Nature 487: 51–6, 2012.

Churchland MM, Shenoy K V. Temporal complexity and heterogeneity of single-neuron activity in premotor and motor cortex. J Neurophysiol 97: 4235–57, 2007.

Churchland MM, Yu BM, Ryu S, Santhanam G, Shenoy K V. Neural variability in premotor cortex provides a signature of motor preparation. J Neurosci 26: 3697–3712, 2006.

Craik K. Theory of the human operator in control systems. I. The operator as an engineering system. Br J Psychol 38: 56–61, 1948.

Desmedt JE, Godaux E. Ballistic contractions in man: characteristic recruitment pattern of single motor units of the tibialis anterior muscle. J Physiol 264: 673–693, 1977. doi:10.1113/jphysiol.1977.sp011689.

Elliott D, Hansen S, Grierson LEM, Lyons J, Bennett SJ, Hayes SJ. Goal-directed aiming: two components but multiple processes. Psychol Bull 136: 1023–1044, 2010.

Elsayed GF, Lara AH, Kaufman MT, Churchland MM, Cunningham JP. Reorganization between preparatory and movement population responses in motor cortex. Nat Commun 7: 1–15, 2016.

Evarts E V. Relation of pyramidal tract activity to force exerted during voluntary movement. J Neurophysiol 31: 14–27, 1968.

Evarts E V, Fromm C. Sensory responses in motor cortex neurons during precise motor control. Neurosci Lett 5: 267–272, 1977.

Evarts E V, Fromm C, Kroller J, Jennings VAA, Kröller J, Jennings VAA. Motor cortex control of finely graded forces. J Neurophysiol 49: 1199–1215, 1983.

Farina D, Fosciand M, Merletti R. Motor unit recruitment strategies investigated by surface EMG variables. J Appl Physiol 92: 235–247, 2002.

Fitts P. The information capacity of the human motor system in controlling the amplitude of movement. J Exp Psychol 47: 381–391, 1954.

Fitts P. Information capacity of discrete motor responses. J Exp Psychol 47: 381–391, 1964.

Flash T, Hogan N. The coordination of arm movements: an experimentally confirmed mathematical model. J Neurosci 5: 1688–1703, 1985.

Freund HJ. Motor unit and muscle activity in voluntary motor control. Physiol Rev 63: 387–436, 1983.

Fu QG, Flament D, Coltz JD, Ebner TJ. Temporal encoding of movement kinematics in the discharge of primate primary motor and premotor neurons. J Neurophysiol 73: 836–854, 1995.

Fu QG, Suarez JI, Ebner TJ. Neuronal specification of direction and distance during reaching movements in the superior precentral premotor area and primary motor cortex of monkeys. J Neurophysiol 70: 2097–2116, 1993.

Georgopoulos AP, Kalaska JF, Caminiti R, Massey JT. On the relations between the direction of two-dimensional arm movements and cell discharge in primate motor cortex. J Neurosci 2: 1527–37, 1982.

Georgopoulos AP, Kalaska JF, Massey JT. Spatial trajectories and reaction times of aimed movements: Effects of practice, uncertainty, and change in target location. J Neurophysiol 46: 725–743, 1981.

Georgopoulos AP, Schwartz AB, Kettner RE. Neuronal population coding of movement direction. Science 233: 1416–1419, 1986.

Griffin DM, Hoffman DS, Strick PL. Corticomotoneuronal cells are “functionally tuned.” Science 350: 667–670, 2015.

Hatsopoulos NG, Xu Q, Amit Y. Encoding of movement fragments in the motor cortex. J Neurosci 27: 5105–14, 2007.

Henneman E, Somjen G, Carpenter D. Functional significance of cell size in spinal motoneurons. J Neurophysiol 28: 560–580, 1965.

Inoue Y, Mao H, Suway SB, Orellana J, Schwartz AB. Decoding arm speed during reaching. Nat Commun 9: 1–14, 2018.

Kalaska JF, Crammond DJ. Cerebral cortical mechanisms of reaching movements. Science (80-) 255: 1517–1523, 1992.

Kalaska JF, Scott SH, Cizek P, Sergio LE. Control of reaching movements. Curr Opin Neurobiol 7: 849–859, 1997.

Karpowicz BM, Ali YH, Wimalasena LN, Sedler AR, Keshtkaran MR, Bodkin K, Ma X, Miller LE, Pandarinath C. Stabilizing brain-computer interfaces through alignment of latent dynamics. bioRxiv 2022.04.06.487388; doi: https://doi.org/10.1101/2022.04.06.487388

Kaufman MT, Churchland AK. Many paths from state to state. Nat Neurosci 19: 1541–1542, 2016.

Ma X, Rizzoglio F, Perreault EJ, Miller LE, Kennedy A. Using adversarial networks to extend brain computer interface decoding accuracy over time [Online]. bioRxiv 2022.08.26.504777; doi: https://doi.org/10.1101/2022.08.26.504777

MacKenzie IS, Buxton W. Extending Fitts’ law to two-dimensional tasks. Conf Hum Factors Comput Syst - Proc 219–226, 1992.

Maynard EM, Hatsopoulos NG, Ojakangas CL, Acuna BD, Sanes JN, Normann RA, Donoghue JP. Neuronal interactions improve cortical population coding of movement direction. J Neurosci 19: 8083–8093, 1999.

Milner-Brown HS, Stein RB, Yemm R. Changes in firing rate of human motor units during linearly changing voluntary contractions. J Physiol 230: 371–390, 1973.

Milner TE. A model for the generation of movements requiring endpoint precision. Neuroscience 49: 487–496, 1992.

Milner TE, Ijaz MM. The effect of accuracy constraints on three-dimensional movement kinematics. Neuroscience 35: 365–374, 1990.

Mollazadeh M, Aggarwal V, Davidson AG, Law AJ, Thakor N V, Schieber MH. Spatiotemporal variation of multiple neurophysiological signals in the primary motor cortex during dexterous reach-to-grasp movements. J Neurosci 31: 15531–43, 2011.

Monk D, Jagacinski R. Fitts’ law in two dimensions with hand and head movements. J. Mot. Behav. : 77–95, 1985.

Moran DW, Schwartz AB. Motor cortical representation of speed and direction during reaching. J Neurophysiol 82: 2676–2692, 1999.

Morasso P. Spatial Control of Arm Movements. Exp Brain Res 42: 223–224, 1981.

Mottet D, Bootsma RJ, Guiard Y, Laurent M. Fitts’ law in two-dimensional task space. Exp Brain Res 100: 144–148, 1994.

Mridha MF, Das SC, Kabir MM, Lima AA, Islam MR, Watanobe Y. Brain-computer interface: advancement and challenges. Sensors 21: 1–46, 2021.

Murata A. Extending effective target width in Fitts’ law to a two-dimensional pointing task. Int J Hum-Comp Interaction 11: 137–152, 1999.

Nicolelis MAL, Ghazanfar AA, Faggin BM, Votaw S, Oliveira LMO. Reconstructing the engram: simultaneous, multisite, many single neuron recordings. Neuron 18: 529–537, 1997.

Novak K, Miller L, Houk J. The use of overlapping submovements in the control of rapid hand movements. Exp Brain Res 144: 351–364, 2002.

Pardoe I. Applied regression modeling. Wiley, 2020.

Port NL, Lee D, Dassonville P, Georgopoulos AP. Manual interception of moving targets I. Performance and movement initiation. Exp Brain Res 116: 406–420, 1997.

Pratt J, Chasteen AL, Abrams RA. Rapid aimed limb movements: age differences and practice effects in component submovements. Psychol Aging 9: 325–334, 1994.

Rouse AG, Schieber MH. Condition-dependent neural dimensions progressively shift during reach to grasp. Physiol Behav 176: 139–148, 2017.

Rouse AG, Schieber MH. Spatiotemporal distribution of location and object effects in primary motor cortex neurons during reach-to-grasp. J Neurosci 36: 10640–10653, 2016.

Rouse AG, Schieber MH. Condition-dependent neural dimensions progressively shift during reach to grasp. Cell Rep 25: 3158–3168, 2018.

Rouse AG, Schieber MH, Sarma S V. Cyclic, condition-independent activity in primary motor cortex predicts corrective movement behavior. eNeuro 9: 1–15, 2022.

Sainburg RL, Ghez C, Kalakanis D. Intersegmental dynamics are controlled by sequential anticipatory, error correction, and postural mechanisms. J Neurophysiol 81: 1045–1056, 1999.

Scott SH. Optimal feedback control and the neural basis of volitional motor control. Nat Rev Neurosci 5: 532–546, 2004.

Soechting JF. Effect of target size on spatial and temporal characteristics of a pointing movement in man. Exp Brain Res 54: 121–132, 1984.

Stein RB, Gossen ER, Jones KE. Neuronal variability: noise or part of the signal? Nat Rev Neurosci 6: 389–397, 2005.

Takeda M, Sato T, Saito H, Iwasaki H, Nambu I, Wada Y. Explanation of Fitts’ law in reaching movement based on human arm dynamics. Sci Rep 9: 1–12, 2019.

Umair A, Ashfaq U, Khan MG. Recent trends, applications, and challenges of brain-computer interfacing (BCI). Int J Intell Syst Appl 9: 58–65, 2017.

Wang W, Chan SS, Heldman DA, Moran DW. Motor cortical representation of position and velocity during reaching. J Neurophysiol 97: 4258–70, 2007.

Woodworth RS. The accuracy of voluntary movement. J Nerv Ment Dis 26: 743–752, 1899.

